# The fidelity of genetic information transfer with aging skeletal muscle segregates according to biological processes

**DOI:** 10.1101/2022.07.18.500243

**Authors:** Sruthi Sivakumar, Ryan William LeFebre, Giulia Menichetti, Hirotaka Iijima, Andrew Mugler, Fabrisia Ambrosio

## Abstract

Maintenance of organismal function requires tightly regulated biomolecular communication. However, with aging, communication deteriorates, thereby disrupting effective information flow. Using information theory applied to skeletal muscle single cell RNA-seq data from young, middle-aged, and aged animals, we quantified the loss of communication efficiency over time. We considered communication channels between transcription factors (TF; ‘input message’) and corresponding target genes (TG; ‘output message’). Mutual information (MI), defined as the information effectively transmitted between TFs and TGs, declined with age. This decline was attributed to escalating biological noise and loss of precision with which TFs regulate TGs (i.e., channel capacity). When we ranked TF:TG pairs by MI, pairs associated with fatty acid oxidation displayed the greatest loss of communication with aging, while the system preserved communication between pairs related to RNA synthesis. These data suggest ineffective communication with aging against a backdrop of resource reallocation to support essential cellular functions.

## INTRODUCTION

All matter undergoes ‘wear and tear’ due to the destructive effects of physical forces; this is a core fundamental truth even for biological systems. The maintenance of proper functioning requires trillions of biomolecules to communicate with each other in an orderly fashion.

However, over time, communication deteriorates, thereby disrupting effective information flow within biomolecular networks (*1*). As a result, essential biological functions are progressively impaired, and organisms become increasingly vulnerable to death (i.e., aging). Aging has been defined as a progressive loss of body function due to the accumulation of damage to molecules, cells, and tissues (*2*).

To date, studies that have estimated escalating biological noise within an aging system have focused on variability in the expression of individual genes and/or signaling pathways (*3–6*). However, such an estimation of biological noise for individual genes does not provide a holistic view of disorder in the aging system. This limitation highlights the need to move beyond isolated molecular snapshots to approaches that account for the complex, dynamic interactions within biological networks. Towards such an integrated approach, in our previous work, we considered the inter-connectivity of biomolecules and quantified network Shannon entropy (*7*). This method enables the examination of molecular relationships and collective behaviors rather than isolated components. We focused on skeletal muscle as a model system given that it has been shown that skeletal muscle health is a good predictor of both morbidity and mortality into older age (*8–10*). We demonstrated that the collective ‘molecular disorderliness’ of the skeletal muscle transcriptome escalates with age (*11*), suggesting that, at the level of the tissue, biomolecular communication becomes increasingly unpredictable over time.

Effective communication is also essential in the regulation of biological processes and function. Cells perform complex tasks by functioning as a network of interacting genes, and successful regulation among genes occurs when the cellular system can produce a reliable and reproducible response to a given external stimulus. In the context of the regulatory network, a communication channel can be defined with transcription factor (TF) expression representing the ‘input message’ and target gene (TG) expression representing the ‘output message’. Effective regulation occurs when the mutual information (MI) between the gene pairs is high, where MI is defined as the information obtained about the input by observing the output. The maximum MI for a given channel over all possible input patterns constitutes the information capacity of the channel (*12*). Although Claude Shannon introduced mathematical concepts to quantify the transfer of information in 1948, it was not until the 2000s that many investigators began to apply these concepts with high rigor to transcriptional regulatory systems (*13–18*). For example, Tkacik et. al. showed that the transcriptional regulatory system can have multiple states carrying more than one bit of information (*13, 18*), thereby generalizing beyond the notion that gene regulatory elements represent a simple “on-off” state. Over the next decade, researchers applied a similar framework to estimate the information transduced by receptors and biochemical signaling pathways (*1, 19–21*). Still unknown, however, is whether and how information flow is altered with aging regulatory networks. To fill this gap, we built upon our previous work in skeletal muscle and here tested the hypothesis that intrinsic biological noise arising from unpredictable fluctuations in gene expression with aging disrupts effective communication between TFs and TGs, leading to compromised function.

## RESULTS

### The amount of useful information transmitted between regulatory gene pairs decreases with age

As a first step to test our hypothesis, we employed an information theoretic approach to quantify the escalating biological noise of the communication network at the level of single cell transcripts. The Tabula Muris Consortium has compiled the transcriptional profiles of multiple tissues at the single-cell level across the lifespan of mice as a part of Tabula Muris Senis (TMS) project (*22*). We used FACS-Smart-seq2 transcript abundance obtained from a total of 3855 cells from the skeletal muscle of young (3 months), middle-aged (18 months), and aged (24 months) mice for these analyses **(Supplemental Figure 1A)**. In agreement with a recent study (*23*), we observed that mean and total overall transcript counts in skeletal muscle decreased with age (**Figure 1A, Supplemental Figure 1B**). To quantify whether molecular disorder of transcript expression within single cells increases with age, we computed Shannon entropy of mean gene counts per animal. A higher Shannon entropy indicates that the probability distribution of gene expression becomes more uniform, thereby decreasing the ability to reliably predict gene expression. Consistent with our previous work using bulk RNA-seq data from skeletal muscle (*11*), we found that entropy at the level of single muscle cells increases in older age (**Figure 1B**).

**Figure 1.**
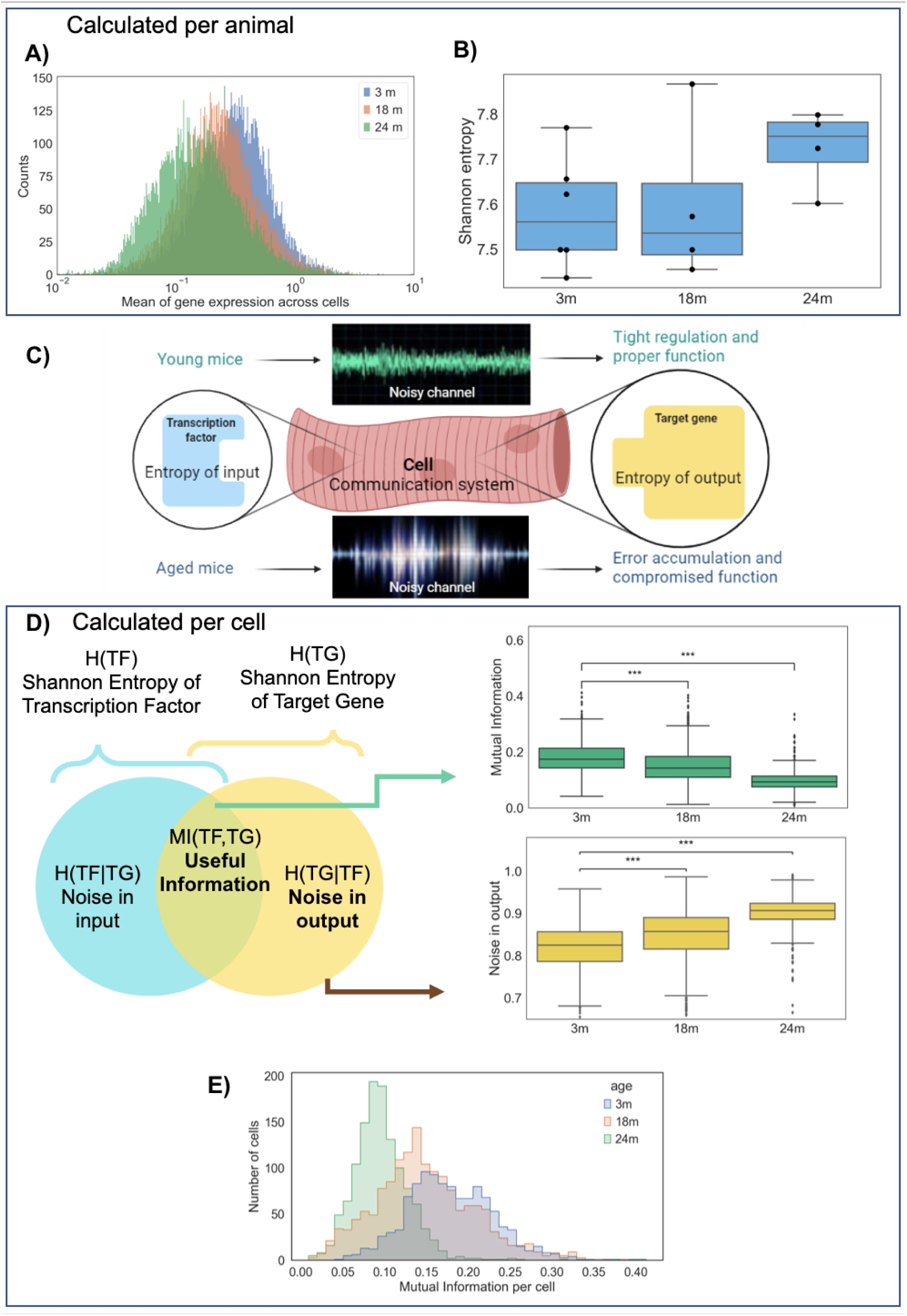
Useful information transmitted in transcriptional regulatory network declines with age as shown by decreasing mutual information (A) Histogram showing mean gene expression across cells decreases with age from single-cell RNA seq Smartseq2 TMS data (Number of cells(n) = 1102, 1521, and 1232 in 3m, 18m, and 24m respectively; male animals: N=4,2,4; female animals: N=2,2,0). The Y-axis indicates the number of genes with that particular mean expression value. (B) Shannon Entropy of mean gene expression per animal increases with age, as per single-cell RNA seq TMS hindlimb muscle data (males: n=4,2,4; females: n=2,2,0). The boxplot with overlaid individual data points indicate median and interquartile range. (C) Schematic representing Shannon’s noisy channel as applied to an aging biological system. We hypothesized that, in an aged system, increasing biological noise leads to impaired communication flow, ultimately contributing to increased error accumulation and compromised function. (D) Venn diagram and schematic showing components of communication channel. Both useful information (mutual information, or MI) and noise in the output (H(TG|TF)) are normalized by mean of entropy of input and output, as described in the methods. The boxplots represent median and interquartile range across age. *** represents p<0.01 for bonferroni post-hoc performed after non-parametric Kruskal Wallis test (Number of cells(n) = 1102, 1521, and 1232 in 3m, 18m, and 24m respectively). (E) Histogram shows that MI declines and that the spread of MI becomes narrow with increasing age.

Next, we wanted to know if the increase in gene expression entropy with aging reflects a decrease in the efficiency of communication between genes. Recent evidence suggests that cell-cell transcriptional ‘noise’, typically defined as the standard deviation of a single transcript, increases with age (*3–6*). Yet, variance in transcript expression as a proxy for noise is limited because it does not give an indication of how effectively the transcripts communicate within the regulatory network. In some cases, greater variance can facilitate better regulation (*1, 24*). To address this limitation, we redefined noise in terms of information lost during communication between regulatory transcripts. Specifically, for a pair of regulatory transcripts, we considered noise as the discrepancy between input and output, which we termed ‘noise in input’ at the regulator end and ‘noise in output’ at the effector end (**Figure 1C, 1D)**.

In our model system, we treated each cell as an individual network of communication channels, with each channel involving an input transcription factor (TF) and an output target gene (TG), we generated a network of TF:TG pairs reported in the TRRUSTv2 database for each single-cell in each age group (*25*). To describe whether and how communication between a given TF and TG pair changes with age, we applied Shannon’s noisy channel coding theorem (**Figure 1C**) (*12*). According to Shannon’s theorem, mutual information (MI) is the amount of information that can be conveyed through a noisy communication channel. We found that average MI, or the average information effectively transmitted between TF:TG pairs, declined with age, as evidenced by a leftward shift in the distribution (**Figures 1D, 1E**).

We next evaluated whether the decline in MI is accompanied by increasing noise in output over time. In the context of our communication channel, output noise refers to factors that collectively disrupt the efficient transfer of information from the input TF to the final output TG. These disruptions can arise from disruption of transcriptional processes or inefficiencies within the regulatory network. Our analysis revealed that output noise progressively increased with age (**Figure 1D**). Of note, we normalized MI and ‘noise in output’ by the mean entropy of input and output to estimate MI, unless otherwise specified (**Supplemental File, equations i-iv**). Our observation of the increased biological noise and a corresponding decrease in MI suggests that TF:TG communication pathways become more unpredictable with age.

### Loss of information with aging is driven by a decreased channel capacity

According to Shannon’s noisy channel theorem, channel capacity (CC) is the maximum rate at which information can be reliably transmitted through a noisy communication channel (*12*). For our system, CC is calculated as the maximum MI possible based on the distribution of TF levels. More simply, CC can be thought of as the maximal precision with which a given TF can regulate its corresponding TG (*13*). CC depends on two quantitative features, range and variability (**Figure 2A)**. Range refers to the difference between the maximum and minimum average output (TG) values observed for a given input (TF) (i.e., the height spanned by the curve in **Figure 2A**). CC is expected to increase with increasing range. Variability refers to the average amount of fluctuation in the output (TG) for a given input (TF), calculated as the variance of the output across all input values (i.e., the average size of the error bars in **Figure 2A**). CC is expected to decrease with increased variability.

**Figure 2.**
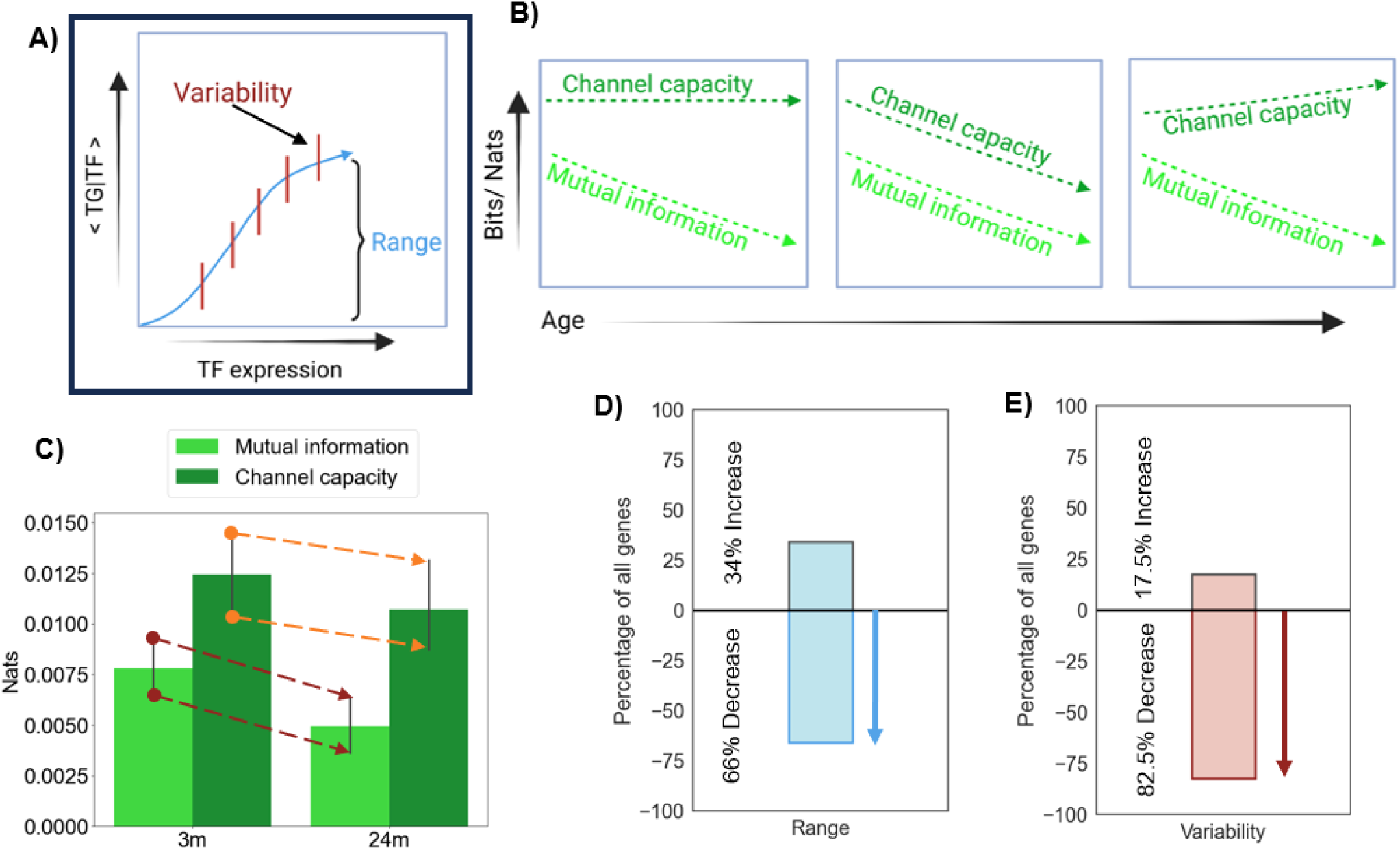
Channel capacity with age declines with age. **(A)** Schematic with hypothetical(dummy) data representing variability and range are two factors that affect the Channel Capacity (CC) value. The conditional probability <TG|TF> on the y-axis is a family of distributions. The plot shows statistics of these family of distribution with respect to TF expression. The blue line represents the conditional averages while the red lines represent the conditional standard deviations. The average of all conditional standard deviations is known as the variability. The difference between maximum and minimum conditional average is the range. The calculations are described in methods. **(B)** Schematic representing hypothesized scenarios for CC changing with age, given that MI declines. **[C, D, E]** Using a coarse-grained four-state model: **(C)** Plot shows that total CC follows mutual information trend, indicating that the precision with which TFs regulate their corresponding TGs decreases over time. The error bar indicates 95% confidence interval. **(D)** Plot shows that range for 66% of TF-TG pairs (n=3021) decreases with age. **(E)** Plot shows that variability for 82.5% of TF-TG pairs (n=3021) decreases with age.

There are three possibilities for how the CC may change relative to the observed decrease in MI with increasing age (**Figure 2B**). One possibility is that CC remains constant over time, meaning that the efficiency of communication decreases even though the system retains its potential for TFs to regulate their TGs with aging. Second, CC may decline with MI, suggesting that the potential of a TF to regulate its TG declines with age, driving a loss of effective communication between the two. Finally, CC could increase over time even as MI decreases.

The interpretation of this third, least likely, scenario would be that the aging system displays an increased regulatory potential, but that other factors preclude the regulatory network from taking advantage of such potential, resulting in faulty communication. We sought to determine which of these possibilities manifests in our aging muscle system.

Computing CC is mathematically challenging, especially for a system such as ours with smooth continuous input and output signals (*13, 21*). Finding the maximum information requires exploring endless possibilities along the smooth input-output curves by brute-force, which is computationally demanding. To simplify, the system was reduced to a basic 4-state model, allowing for easier calculations using straightforward formulas **(Supplemental Figure 2A, 2B)**. The 4 states we defined were based on simplifying input-output expression levels into binary variables: “0” if the gene count is zero and “1” if the gene count is non-zero. Hence, the 4 states for TF:TG mappings were (0, 0), (0, 1) (1, 0), and (1, 1). Such binary reduction is common for RNA-seq data, particularly because many of the outputs are zero(*26*). Consistent with our findings when fine-grained continuous binning was used **(Figure 1D)**, we observed an overall decreasing trend for MI with age using a coarse-grained model (**Figure 2C**). Furthermore, we found that this decrease was accompanied by a reduced CC (**Figure 2C**). We also confirmed that this finding using a four-state model is not due to reduced gene expression at binary levels by computing MI and CC for a range of nonzero thresholds of TF and TG expression **(Supplemental Figure 3)**. These results suggest that the loss of effective communication over time is likely driven by a gradual decrease in the precision with which TFs can regulate their TGs.

The next logical question was, *what causes CC to decline with age?* As described above, range and variance are the two factors that contribute to our calculations of CC. We found that, with aging, most TF:TG pairs displayed a decrease in range and variance (**Figure 2D, E**). This implies that the TG expression becomes more uniform across cells (i.e., lower variance), but with less influence from TFs (i.e., reduced range). The reduced variance is to be expected because, for any system, variance becomes constrained as gene expression values approach zero (**Supplemental File, under “Range and variability as they change with channel capacity”**).

### Transcriptional regulation is preferentially preserved while fatty acid metabolism is compromised with aging

The above findings demonstrate that an aging skeletal muscle system displays an overall decrease in the transmission of ‘useful’ information, which is due to the inability of TFs to effectively regulate TGs within the regulatory network. To better understand how gene regulation may influence the escalating disorder with aging, we calculated the total Shannon entropy of genes in the transcriptional regulatory network, as reported in the TRRUSTv2 database, and compared it to the entropy of genes not in the database. Whereas the total entropy of genes not in the database increased over time (**Supplemental Figure 4A)**, TFs and TGs within the transcriptional regulatory network displayed minimal change in entropy with aging (**Supplemental Figure 4B, C**). This raised the interesting possibility that regulation renders genes less susceptible to molecular disorder as the organism ages.

The above observation led us to ask whether there may be a distinct set of genes within the transcriptional regulatory network that are preferentially regulated according to biological relevance. To answer this question, we generated a list of TF:TG pairs and their corresponding MI values across young and aged groups using continuous binning. TF:TG pairs were then divided into two categories: one subset of genes that displays a *preserved* MI with age, and a second subset that displays a *compromised* MI with age. For this, the gene subset with preserved MI was defined *a priori* as starting with a high MI (>0.3) in young and persisting over time into old age. In contrast, the gene subset that displayed a compromised MI was defined as starting with a high MI (>0.3) in young but decreasing over time. According to these criteria, 248 gene pairs were in the top 20^th^ percentile for preservation of MI with aging, and 248 gene pairs displayed the greatest extent of compromised MI (80^th^ percentile; **Figure 3A**). As expected, there was minimal change in the MI and CC for preserved genes over time (**Figure 3B**), while both MI and CC were significantly decreased for compromised genes (**Figures 3C**).

**Figure 3.**
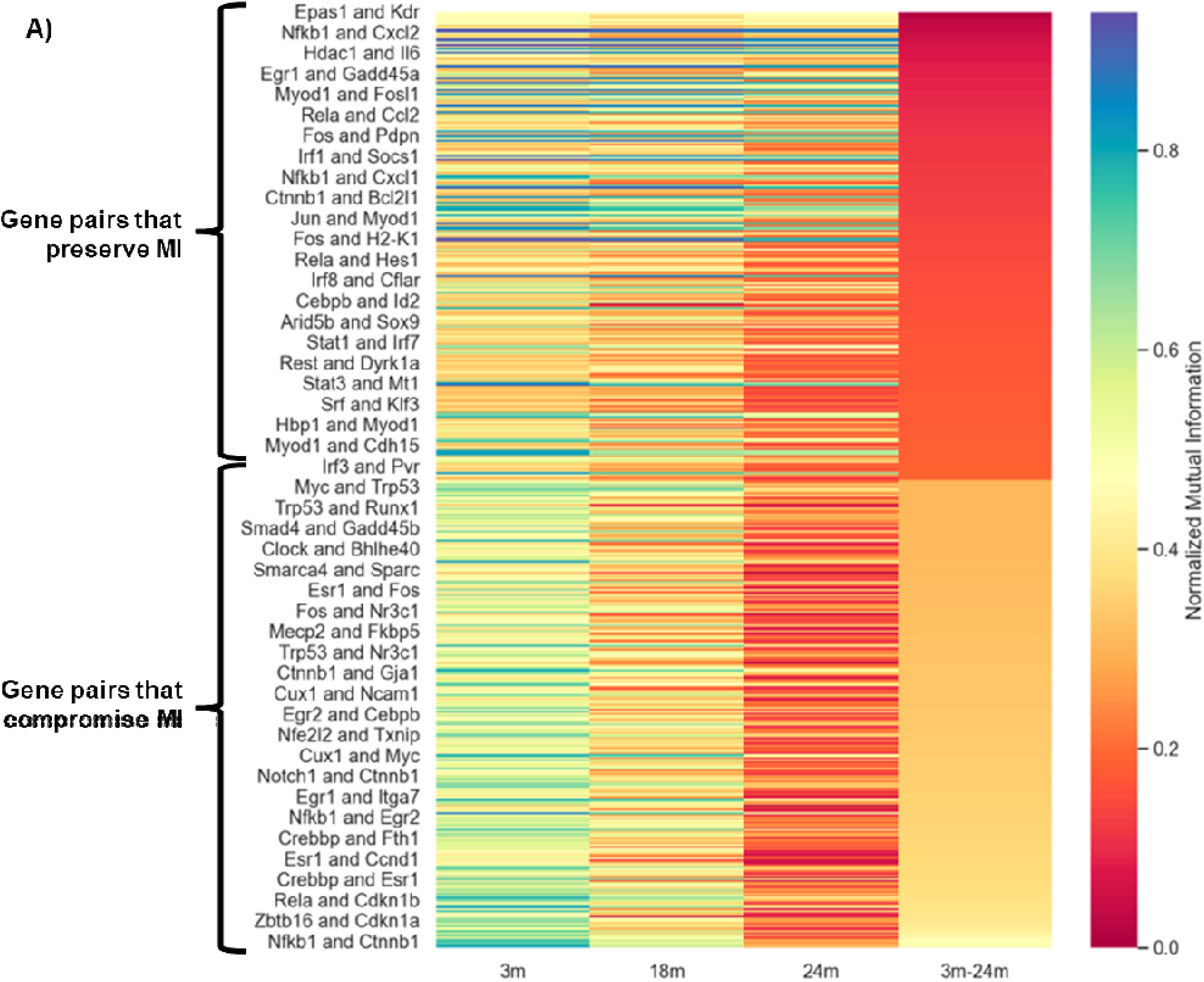
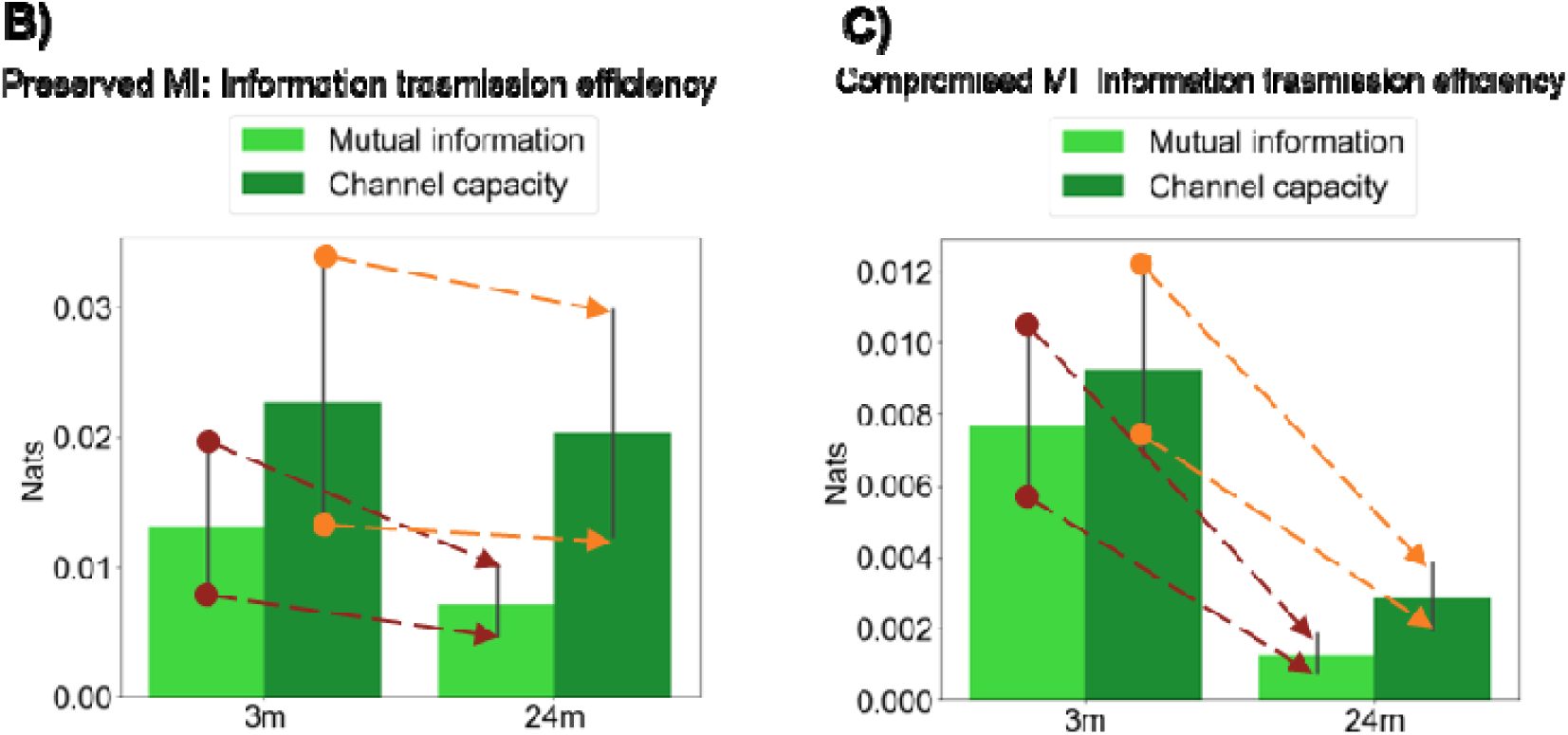
TF:TG pairs can be segregated into a sub-set that display preserved information flow and channel capacity versus a sub-set in which information transfer is compromised. **(A)** The heat map represents mutual information for gene pairs per age group calculated with fine-grained continuous binning, as described in methods. The gene pairs are arranged in decreasing order of difference in MI from 3m-24m. Genes that preserve MI have >0.3 normalized mutual information when young and are in the top 20th percentile of values when considering the difference between 3m-24m. Genes that are defined as displaying compromised MI have >0.3 normalized mutual information when young but are in the bottom 20th percentile when considering the difference between 3m-24m (i.e., these genes displayed the greatest loss of MI over time). The colorbar indicates the value of normalized mutual information (between 0 and 1), with equations are described in the methods. **[B-C]**: We used a four-state coarse grained model to obtain the following results. **(B)** Channel capacity for preserved genes follows MI trend, indicating that the maximal precision with which a given TF can regulate its corresponding TG remains relatively steady with age. The error bar indicates 95% confidence interval. Mann Whitney U test was performed. There was no significant change in MI or CC. **(C)** Channel capacity for compromised genes also follows mutual information trend indicating that the maximal precision with which a given TF can regulate its corresponding TG declines steeply with age.The error bar indicates 95% confidence interval. Mann Whitney U test was performed. MI and CC sign ficantly decreased with aging (*** p<0.01).

Understanding the characteristics of regulatory networks that either remain resilient (preserved pairs) or fail (compromised pairs) with aging has the potential to reveal valuable insights into how the system re-allocates resources and adapts over time. Therefore, we next characterized the biological function of the TGs that displayed preserved versus compromised communication with age. To characterize the function of these genes that are possibly working together, we created a Weighted Gene Co-expression Network (WGCNA) using bulk RNA-seq skeletal muscle data from 54 mice across the lifespan (1 month-27 months) in the Tabula Muris Senis database (*27, 28*) **(Figure 4A)**. From the aging bulk RNA-seq data, we identified 16 such highly co-expressed modules. Out of the 16 modules, the 14^th^ module was highly enriched for the TGs that displayed preserved communication over time whereas the 16^th^ module was enriched for TGs displaying compromised communication over time **(Figure 4B)**. We then used EnrichR over-representation analysis followed by REVIGO to summarize the biological processes associated with these modules. The most highly represented module for preserved genes was predominantly linked to RNA synthesis and processing, whereas the dominant module for compromised genes was overwhelmingly linked to fatty acid metabolism **(Figure 4C)**.

**Figure 4.**
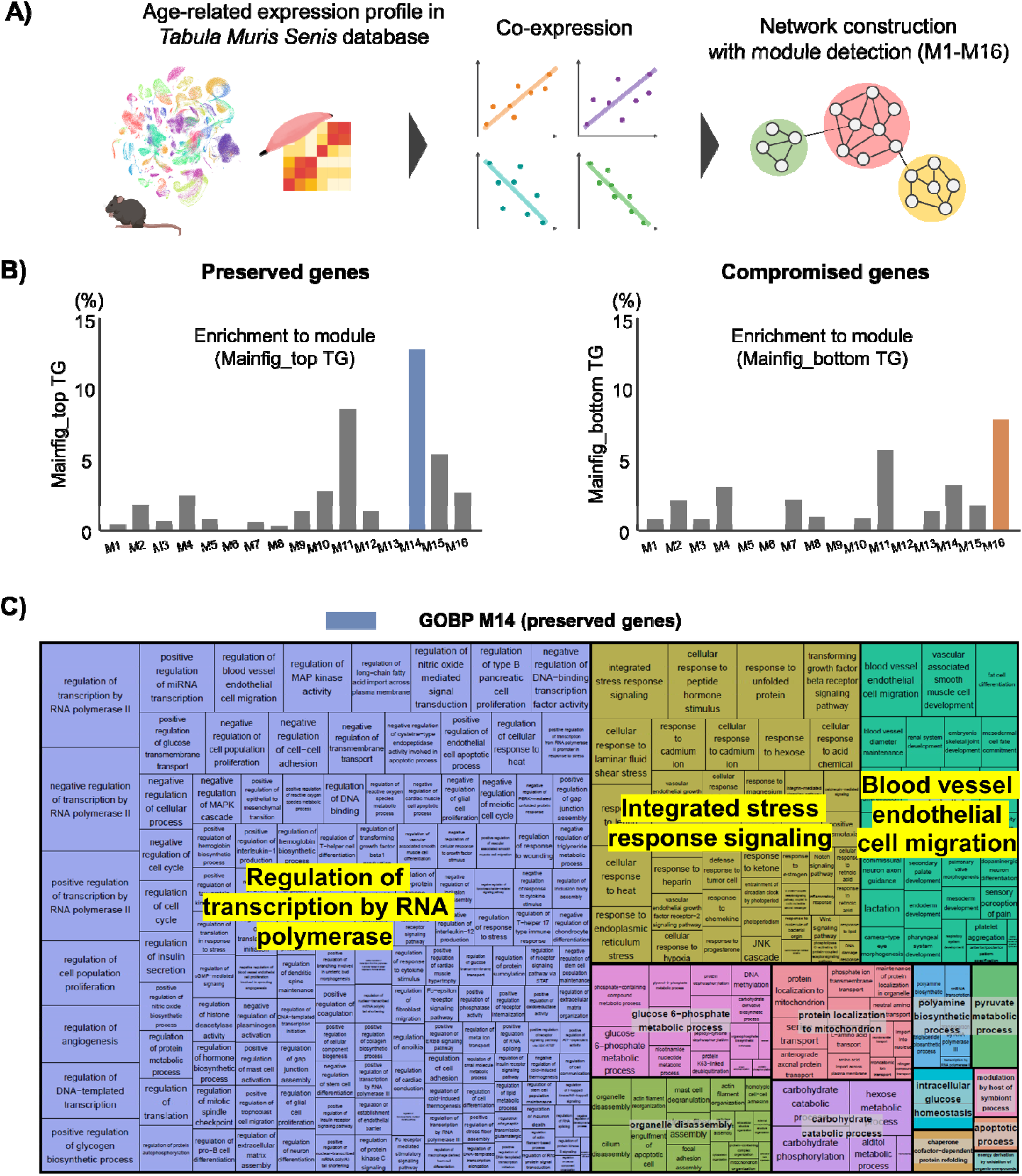
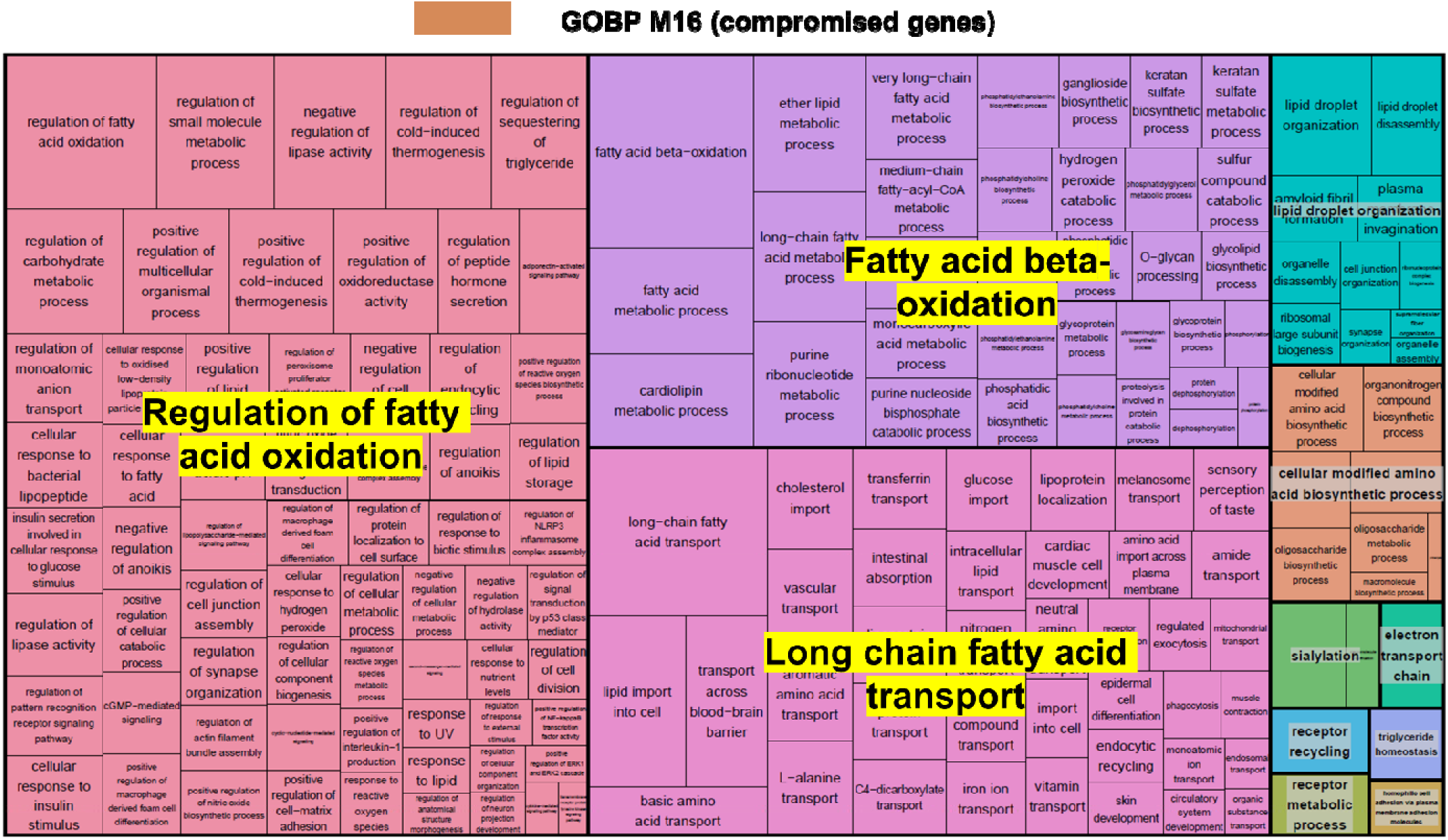
Preserved and compromised genes displayed distinct enrichment to functional module. **(A)** Bulk RNA-seq of skeletal muscle from 54 mice across the lifespan (ages 1 month – 27 months) in the Tabula Muris Senis dataset was used to construct weighted gene co-expression network that reflects age-related biological changes in murine skeletal muscle. Sixteen key modules were detected based on soft thresholding specified in the methods. **(B)** Module 14 was highly enriched for TGs that preserve MI, and module 16 was highly enriched for TGs that compromise MI. **(C)** Gene ontology (GO) enrichment using EnrichR with subsequent REVIGO revealed that Module 14 was primarily associated with regulation of transcription (large blue rectangle) whereas Module 16 was primarily associated with fatty acid metabolism (large red rectangle). Each rectangle represents a supercluster GO, visualized with different colors. The size of the rectangles was adjusted to reflect the p value of the GO term calculated by Top GO (ie, the larger the rectangle, the more significant the GO term).

We also validated these findings using a second, independent dataset that also used FACS to isolate muscle cells followed by Smartseq2 sequencing with cells from young (*n* = 253) and old (*n* = 124) mice (*3*). As observed using the TMS dataset, we found that entropy generally increased, MI across TF:TG pairs decreased, and noise in output (TG) increased with age (**Supplemental Figure 5A-C**). This result suggests that the loss of information with age is a robust feature whose causes include, but are not limited to, a global decrease in expression levels. Moreover, similar to our observation with TMS (**Figure 4**), module 14 representing RNA synthesis and processing was the most enriched for preserved TGs, while module 16 representing fatty acid oxidation was the most enriched for compromised TGs (**Supplemental Figure 5D, 5E**).

Taken together, the enrichment of the preserved communication module in processes related to RNA synthesis and processing suggests that the aging skeletal muscle system prioritizes resources associated with maintenance of basic cellular functions, such as gene expression and protein production. However, the enrichment of the compromised communication module in processes related to fatty acid metabolism may reflect a reduction in the ability of muscle to efficiently manage metabolic resources.

## DISCUSSION

While a mathematical theory of communication was first proposed over 70 years ago, its application to quantify transcriptional information processing with age, as we have done here, is novel. Our information theory-based approach offers an unbiased method to quantify the loss of efficient biomolecular communication with aging at the resolution of single cells (*29–31*). Using skeletal muscle as a model, we found that the amount of useful information transmitted between a TF and its TG progressively declines over time. Further, we found that the precision with which TFs can regulate their TGs decreases with age, and that this loss of regulatory capacity may be a fundamental driver of deteriorating biomolecular communication. Finally, by classifying gene pairs into preserved and compromised sets, we discovered that functions relating to transcriptional regulation were preferentially maintained, while functions related to fatty acid metabolism declined, highlighting the shifting biological priorities of the system as it ages.

From a biological perspective, our information theoretic approach offers the advantage of disentangling unique components of transcriptional heterogeneity and evaluating how this heterogeneity evolves with aging. Whereas previous studies have demonstrated that variability in gene expression, or transcriptional heterogeneity, increases with time (*3–5*), few studies have investigated distinct aspects of transcriptional regulation that contribute to the overall increased heterogeneity with aging. Perez-Gomez et al. referred to the random, or stochastic, transcriptional variability as “transcriptional noise”, while non-random transcriptional variability, which may result from deterioration of systemic regulation, was defined as “transcriptional drift” (*32*). In their work, Perez-Gomez et al. noted that a major challenge lies in distinguishing changes in which the expression of a single gene drifts away from its original level due to a lack of regulation versus adaptive changes that are initiated by regulated processes (*32*). The authors also point out that this distinction can be better made when correlation or co-expression of multiple genes is taken into consideration. In line with this proposition, we defined noise with respect to a pair of genes in a communication channel, and we focused on how efficiency of transcriptional regulation changes with age. Given that MI reflects non-random transcriptional variability, our results suggest that preserved genes display less transcriptional drift while compromised genes display increased transcriptional drift with increasing age. It is important to note the difference in interpretation of variance versus MI. MI is calculated using the joint distribution of input TF and output TG, thereby estimating the association between the entire gene expression distribution of a given TF and TG pair. Thus, MI could decrease because the variance of either the TF or TG goes up with age, but it could also go down because the range of expression levels goes down with age; both effects are captured. In this way, MI is a more agnostic, comprehensive, and interpretable measure to quantify how information flow changes with age as compared to more traditional measures of variance alone.

From a mathematical perspective, our information theoretic approach has several advantages over the conventional statistical measures such as covariance, correlation coefficient, or linear regression. Although these latter measures are typically employed to describe variability/heterogeneity in an aging system (*3–5*), the use of MI as first defined by Shannon has several advantages in the context of a biological system (*12, 16*). First, MI is assumption-free when considering the mathematical function between TFs and TGs. That is, unlike linear regression, there is no linearity assumption, thereby allowing us to consider all possible non-linear dependence that is typical of a complex system. In addition, MI is applicable to both continuous and discrete variables, and it is invariant to reparameterization. Stated otherwise, unlike the Spearman correlation coefficient or covariance, the MI between raw counts of TF and TG is the same as the MI between any one-to-one function of TF and TG, e.g., MI (TF; TG) = MI(log_2_(TF); log_2_(TG)). This is particularly valuable given that we used normalized scRNA-seq gene counts. Another advantage of our approach is that MI also obeys the data processing inequality, where information in the output relative to the input is necessarily either lost or stays the same at each noisy step in the transmission process but is never “spontaneously” created(*33*). Finally, MI has a clear quantitative interpretation as information, i.e., there would be 2^MI(TF;TG)^ distinguishable levels of TG for the range of TF expression for MI measured in bits (*16, 33*).

Since the numerical value of MI will remain invariant to the above effects, usage of MI enhances the reproducibility and rigor of our studies.

Given these advantages of an unbiased information theoretic approach, the next critical consideration is to place the observed decline in MI into a biologically relevant context. The observed decline in CC with age suggests that TFs lose the ability to precisely control their respective TGs. While we obtained well-established known TF:TG associations from the TRRUSTv2 database, we did not directly evaluate cellular conditions that may contribute to this loss of control. Biological changes resulting in CC decline may include, for example, epigenetic alterations that affect TF binding, resource re-allocation, waste accumulation, etc. Indeed, our data suggest that resource allocation may not be the same for different biological functions as the organism ages. Specifically, genes associated with RNA transcription and processing displayed a preserved MI over time, whereas genes associated with fatty acid oxidation displayed the greatest loss of MI. This finding suggests that the organism achieves preferential preservation of cellular functions by regulating certain gene pairs more tightly than others. We note that the model developed here does not consider mechanistic biological parameters, such as chromatin accessibility, histone modifications, or post-translational modifications since scRNA-seq data is not enough to capture these effects. While it would be interesting to better understand how these factors contribute to the loss of biomolecular communication with aging, such analyses would require evaluation of these features together with transcriptomic profiles in the same cell— something that is not technically possible at this time.

Waddington suggested that a robustness to perturbation reflects a long-term prioritization for optimal phenotypes(*34*). Broadly speaking, canalization is an evolutionary process in which the final phenotype persists even in the face of challenges or perturbations(*35*). For example, genetic canalization is when the phenotype remains stable in face of genetic mutations or variations. Environmental canalization is when the phenotype remains stable in response to environmental variations, such as temperature, pressure, etc. In our case, we found that preserved genes remain stable in the presence of age-related perturbations, a phenomenon we termed “age-based canalization”. It has been suggested that canalization is an inevitable consequence of the evolution of complex systems that is designed to increase chances of survival amid complex dynamic processes (*34–37*). Specifically, in a setting of finite resources of an aging organism, the highly optimized complex transcription network may undergo trade-offs that prioritize robustness at the expense of resource use, performance, or control (*37, 38*). This is consistent with our observation of preserved expression of genes associated with transcriptional regulation at the expense of genes associated with fatty acid metabolism in aging skeletal muscle. The loss of communication in TGs associated with fatty acid oxidation is in line findings showing a shift in metabolism with age-related muscle loss (i.e., sarcopenia)(*39–41*). Although it is known that fatty acid oxidation can be manipulated to improve aging skeletal muscle phenotype, further mechanistic studies are warranted to understand molecular interplay underlying these effects.

Although there are several advantages of this approach, limitations should be noted. First, we used gene expression values from single-cell RNA seq data, which contains a combination of technical and biological variability. Distinguishing between these two types of variability is challenging. Since MI is invariant to reparameterization, the model captures core properties of biology despite performing transformations and normalization to the raw transcript counts. Here, we have used FACS-Smartseq2 data, which offers higher gene coverage and relatively less sparse data as compared to the more popular 10X Chromium(*42, 43*). However, FACS isolation may stress cells and affect gene expression profile. As such, our approach will become more accurate as single cell sequencing technologies become less stressful on cells, provide higher gene coverage, and produce less sparse data. We also note that, to increase statistical power, the sample sizes per age group need to be increased and balanced according to sex in future studies. Finally, it would be interesting in future studies to design an *in vivo* experiment to measure time-series measurement of transcript expression levels as the organism ages(*44*). This can potentially serve as a validation of our current findings about preferential preservation of biological functions.

Taken together, the application of an information theoretic approach serves as a biologically intuitive mathematical model that has potential to describe the progressively disrupted cellular communication with age. Our findings demonstrate that increasing biological noise can be quantified using the mathematical basis of communication and suggest that this increase in noise may be preferentially regulated depending on the gene function. Safeguarding of specific pathways into old age contrasts with the popular stochastic error accumulation theory of aging, which purports that molecular mistakes are random (*45*). An enhanced understanding of how certain biological functions are given priority (i.e., ‘biological wisdom’) and the consequence of such trade-offs, may aid in the future design of targeted therapeutics with the long-term goal of sustaining organismal health and function over time.

## MATERIALS AND METHODS

### Data pre-processing

We accessed Tabula Muris Senis (TMS) FACS-Smartseq2 single-cell RNA seq dataset for analyses, <u>DOI:</u> https://doi.org/10.6084/m9.figshare.12654728.v1(21). Since we used skeletal muscle as our model system, we used processed h5ad files obtained from limb muscle across the lifespan of mice (3 months, 18 months, and 24 months). The raw data and steps for the preprocessing are described along with the code in the Github repository, https://github.com/czbiohub-sf/tabula-muris-senis/tree/master. To compare the global transcriptional landscape with aging, we analyzed high resolution full-length transcript data which was generated using FACS-isolation followed by Smart-seq2 sequencing. We discarded very lowly expressed genes for our analyses by only selecting genes that were expressed at non-zero levels in at least 10 cells per age group. Then we took the intersection of genes, across age groups, that passed this filter. Next, to create a transcriptional regulatory network for single cells, we accessed the TRRUSTv2 database (https://www.grnpedia.org/trrust/), which has documented text-mining based transcriptional regulation for mouse and human genes (*24*). We created edge lists for all cells in each age group based on reported TF:TG interactions for mice. After mapping to the regulatory network, there were a total of 1422 genes. Scanpy and anndata were used for scRNA-seq analyses and preprocessing.

### Study Rigor

The TMS dataset is comprised of different sample sizes of male and female animals across the age groups, namely, 2 females and 4 males in young (3 months), 2 females and 2 males in middle-aged (18 months), and 4 males in aged mice (24 months). The total number of cells analyzed from young, middle-aged, and aged were 1102, 1521, and 1232 respectively. UMAP plots provided clusters based on cell type in **Supplemental Figure 1B**. All cell types were present in both male and female animals. However, we were unable to test the effects of sex on information flow calculations because we are limited by low sample numbers for male and female in each age group. For all subsequent analyses, we included all cells from both sexes. The TMS single-cell RNA seq source data is open-access and all the analyses files are documented on Github for reproducibility.

### Information theory computation

Python packages were used for single-cell data wrangling, information-theory based interrogation and visualizations. Shannon entropy and normalized mutual information functions from scikit-learn package were used in **Figure 1**. Specifically, for **Figure 1A**, we calculated and plotted the mean gene counts per cell on a logarithmic scale. For **Figure 1B**, we calculated the mean gene counts per animal and performed logarithmic binning on these mean values, similar to **Figure 1A**, and then calculated Shannon entropy of the binned value. Conditional entropy (‘noise in output’) was calculated as the difference between entropy and normalized mutual information score functions from scipy.stats v1.10.0 and scikit-learn v1.2.2 Python packages, respectively. Normalized Mutual Information (‘normalized_mutual_info_score’ function, referred to as “MI” throughout) is a normalization of the MI score to scale the results between 0 (no mutual information) and 1 (perfect correlation). We note that for this function, each gene expression value was considered its own bin (or “continuous binning”) for the calculation of MI. For **Figure 1D and 1E**, we computed MI of all TFs and corresponding TGs in a given cell and then presented the MI per cell. In **Figure 3A**, we used the same continuous binning MI function, but computed MI for a particular TF and TG (each row in the heatmap) across all cells in each age group. The normalization factor for both MI and noise in output is the mean entropies of input and output. Additionally, derivations for four state-model computations of mutual information and channel capacity used in **Figure 2** are described in **Supplemental File**. In the following equations, TF is the gene expression value for the transcription factor (input), TG is the gene expression value for the target gene (output), N is the length of the list of expression values. H(TF) and H(TG) are Shannon entropy of TF and TG, respectively, and P represents probability of expression value for TF or TG.

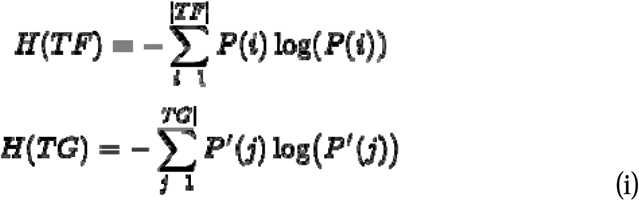

Mutual Information between TF and TG given by MI (TF, TG) is then given by,

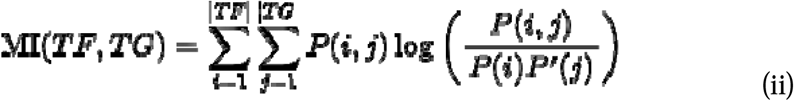

The probability P(i,j) can be written in terms of intersection of TF and TG expression values.

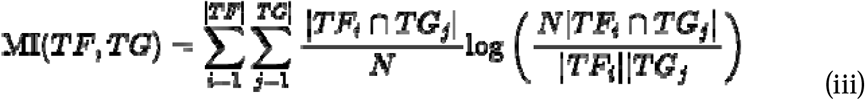

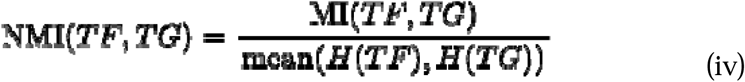

Matplotlib and seaborn packages were used to make histograms, line plots, heatmap, and Venn diagrams. BioRender was used to make the schematic (**Figure 1C**). Matplotlib and seaborn packages were used to make histograms, line plots, heatmap, and Venn diagrams for all figures.

### Data-driven network construction via WGCNA

This study used the WGCNA package to build a weighted gene co-expression network using the the archived bulk RNA-seq data from Tabula Muris Senis of murine skeletal muscle across the lifespan (1, 4, 6, 9, 12, 15, 18, 21, 24, and 27 month old; n = 54 samples in total)(*46*). Raw count data was normalized by count per million (CPM), filterByExpr function, and Trimmed Mean Mvalue (TMM) using R/Bioconductor package edgeR with default parameters(*47*), which finally yielded 8,389 genes. The key parameter, β, for weighted network construction was optimized to maintain both the scale-free topology and sufficient node connectivity as recommended in the manual. A topological overlap matrix (TOM) was then formulated based on the adjacency values to calculate the corresponding dissimilarity (1-TOM) values. Module identification was accomplished with the dynamic tree cut method by hierarchically clustering genes using 1-TOM as the distance measure with a minimum size cutoff of 30 and a deep split value of 2 for the resulting dendrogram. A module preservation function was used to verify the stability of the identified modules by calculating module preservation and quality statistics in the WGCNA package(*28*).

### Functional characterization of transcriptome using pathway enrichment analysis

To determine the biological function of genes of interests, gene ontology (GO) enrichment analysis was performed by Enrichr software(*48*). Subsequently, REVIGO software was applied to summarize redundant GO terms and visualize the summarized results via treemap(*49*).

## Acknowledgments

This work was funded by NIA RO1 AG052978 (FA), R01 AG061005 (FA), and R01AG082739 (FA and AM). We thank Center for Research and Computing at the University of Pittsburgh that provided resources for providing the platform for analyses. We also thank the attendees of Gordon conference on Stochastic Physics in Biology (2021), Gordon conference on Systems Aging (2022), and Keystone conference on Single-cell biology (2022) for engaging in fruitful conversations.

## Author contributions

Conceptualization: SS, FA

Methodology: SS, FA, AM, RWL

Investigation: SS, RWL, HI

Visualization: SS, RWL

Funding acquisition: FA

Project administration: FA

Supervision: FA, AM

Writing – original draft: SS, FA

Writing – review & editing: SS, FA, AM, RWL, HI, GM

## Data and materials availability

All files and code to reproduce all figures are available on Github (https://github.com/sruthi-hub/Aging_mutual_info_TMS_FACS). Raw data and preprocessing code are available here: https://github.com/czbiohub-sf/tabula-muris-senis/tree/master.

## Supplemental Figures

**Supplemental Figure 1:**
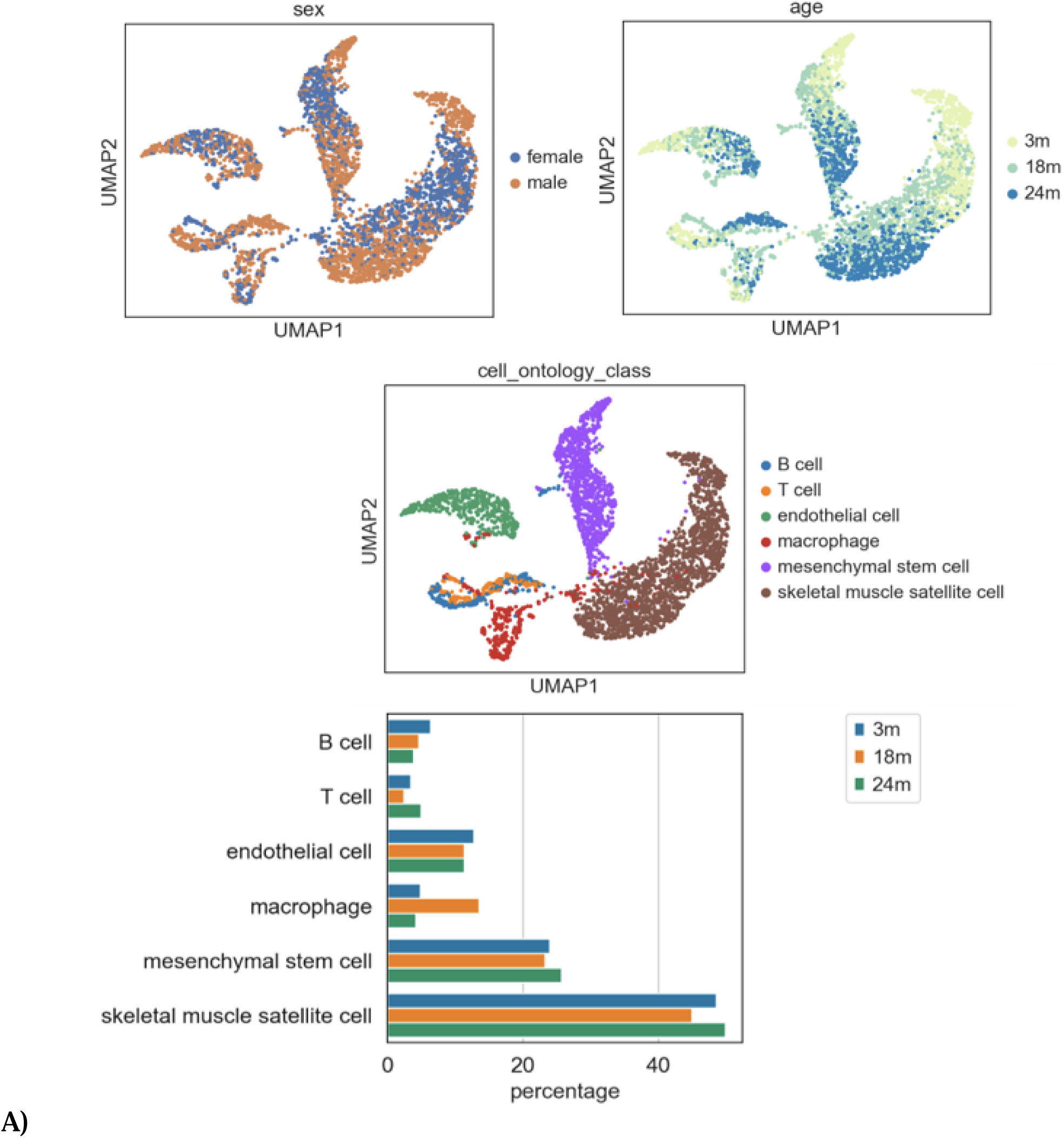
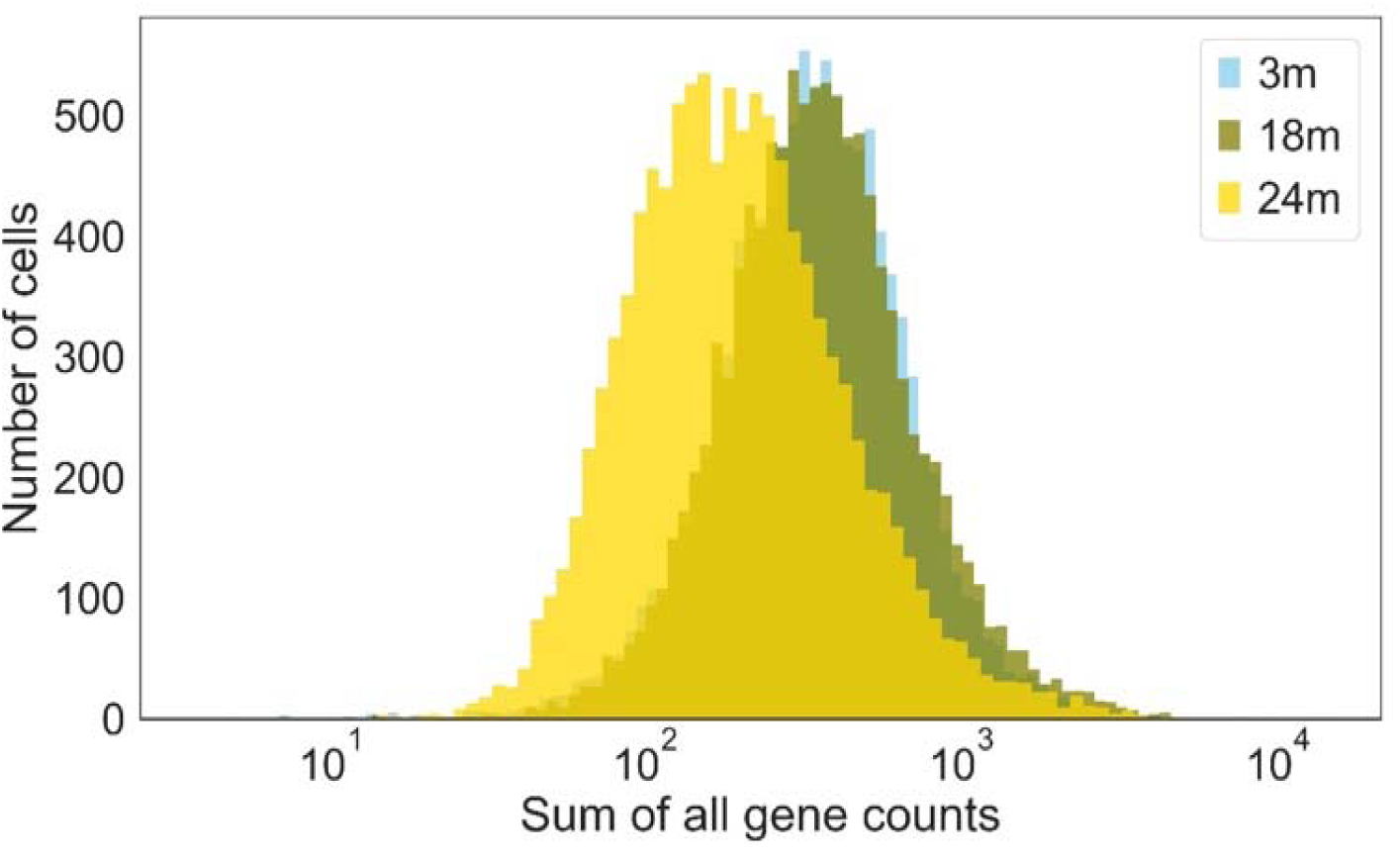
Overview of TMS Smartseq2 limb muscle data. **(A)** First three panels show UMAP embeddings with sex, age and cell type for the TMS FACS-Smartseq2 data that was used for the main figures. The bar plot shows the cell type percentage distribution with age (Number of cells(n) = 1102, 1521, and 1232 in 3m, 18m, and 24m respectively). **(B)** The sum of all gene counts per cell for TMS FACS-Smartseq2 limb muscle declines with age.

**Supplemental Figure 2:**
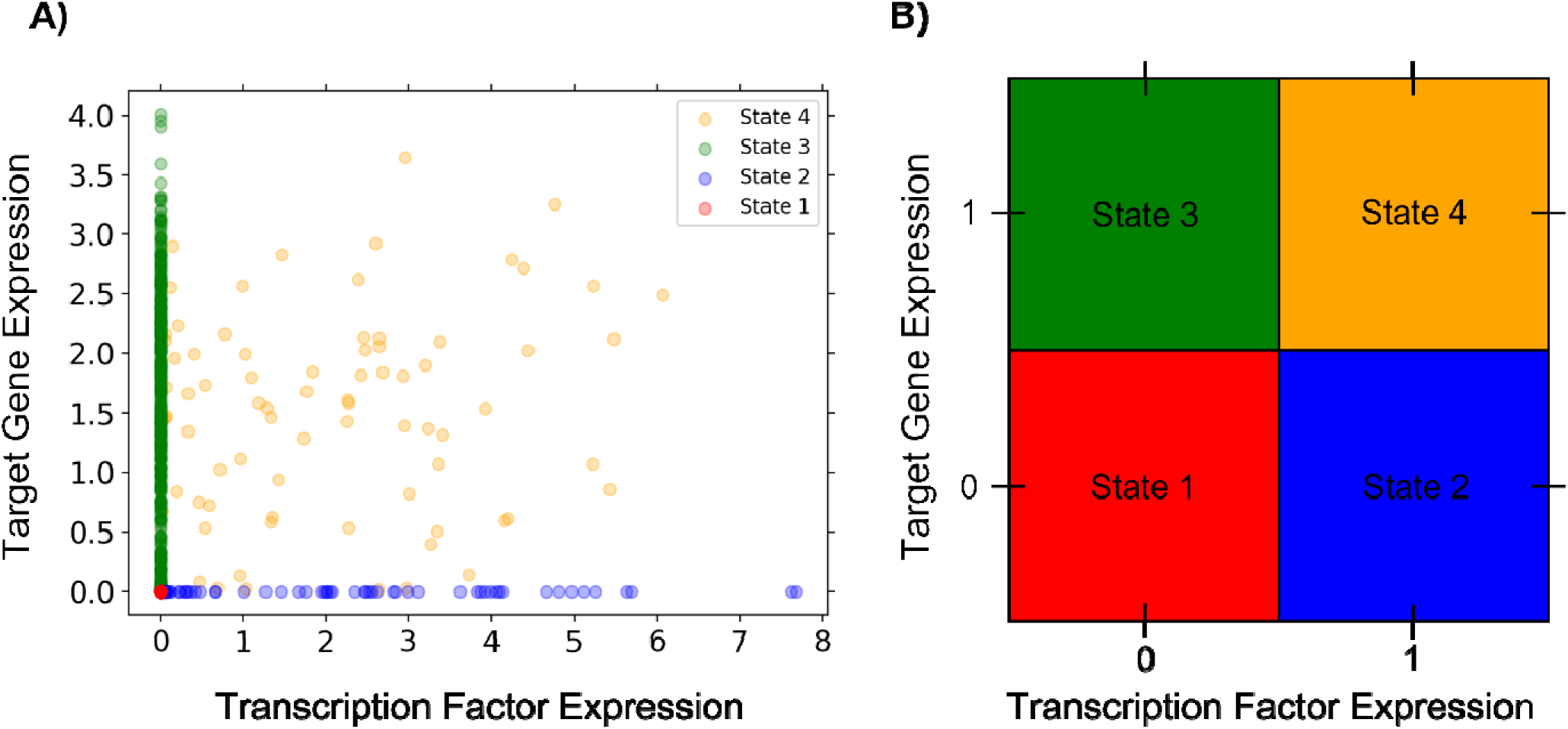
Coarse-graining data into four states. **(A)** Example of a TF-TG pair exp levels colored in four states. **(B)** The four states into which the data are coarsened.

**Supplemental Figure 3:**
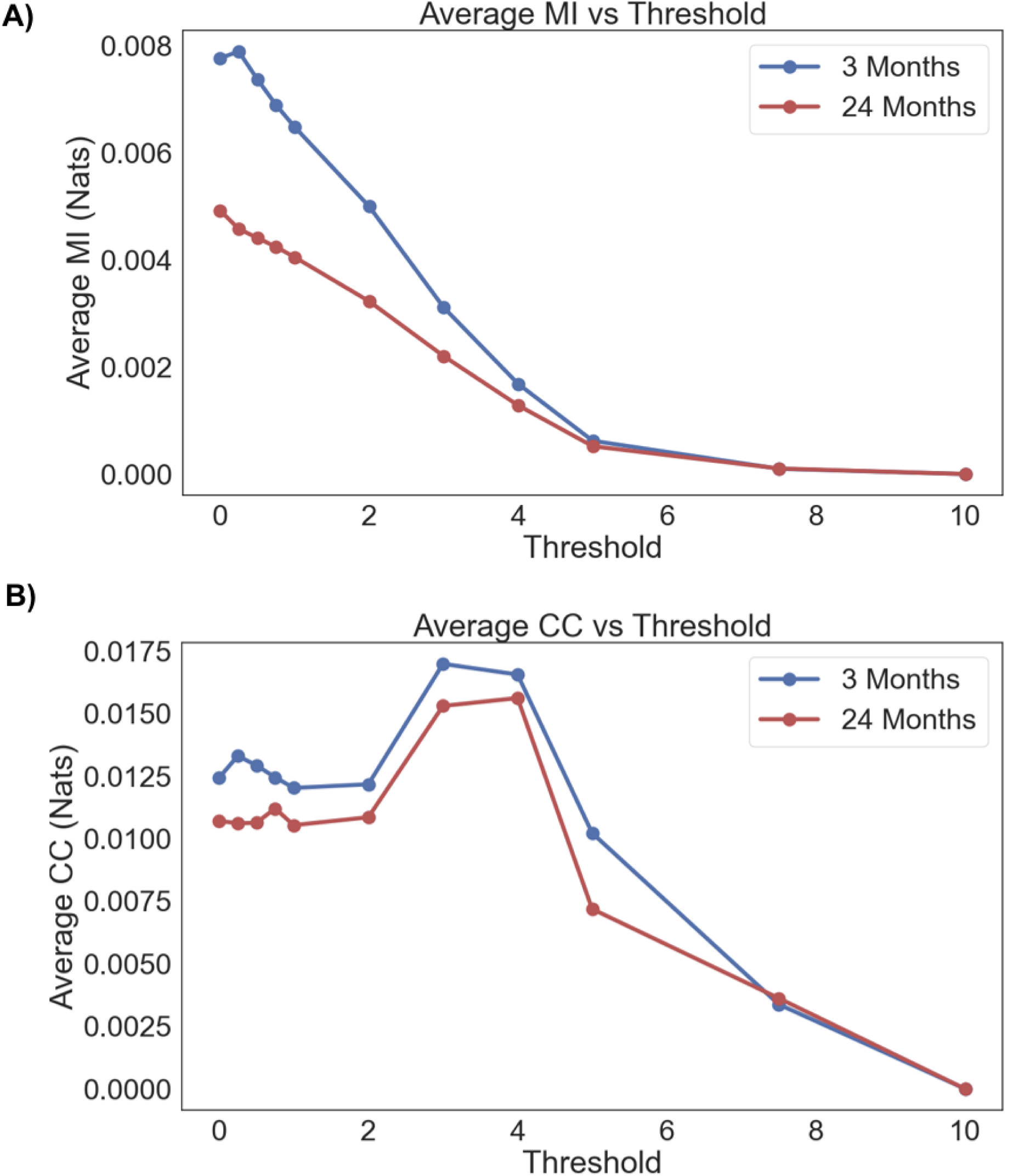
The qualitative trends of mutual information decreasing with age and the channel capacity in the four-state model are robust to non-zero thresholding. **(A)** Average mutual information across gene pairs vs non-zero threshold level. For reasonable non-zero threshold values, the mutual information decreases with age from 3 months to 24 months until all curves converge as the threshold gets too large. **(B)** Average channel capacity across gene pairs vs non-zero threshold level. Channel capacity follows a similar trend to mutual information with respect to age. For reasonable threshold values, channel capacity decreases from 3 months to 24 months before they converge.

**Supplemental Figure 4:**
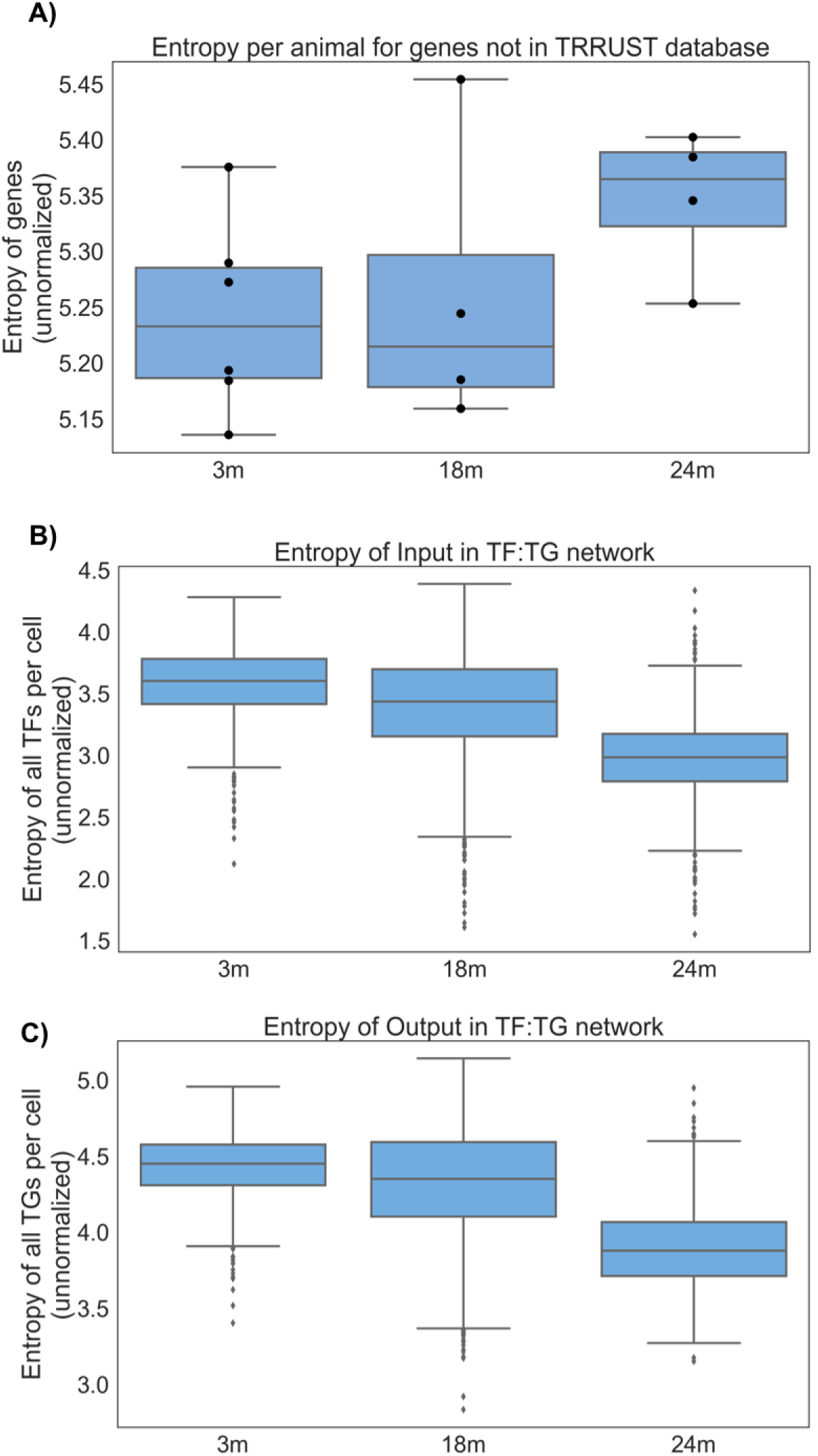
Shannon entropy of genes without normalization. **(A)** Shannon entropy of genes that are present in TMS Smartseq2 limb muscle data but that not a part of the TRRUST database increases with age. Entropy is calculated per animal, similar to Figure 1B (Number of cells(n) = 1102, 1521, and 1232 in 3m, 18m, and 24m respectively; male animals: N=4,2,4; female animals: N=2,2,0). The boxplot with overlaid individual data points indicates median and interquartile range. The increase is not statistically significant. **(B)** Shannon entropy of genes that are a part of the transcriptional regulatory network, i.e., transcription factors, decreases monotonically with age. This is represented by the full blue circle in the Venn Diagram (H(TF) in Figure 1D). **(C)** Shannon entropy of genes that are a part of the transcriptional regulatory network, i.e., target genes, decreases monotonically with age. This is represented by the full yellow circle in the Venn Diagram (H(TG) in Figure 1D).

**Supplemental Figure 5:**
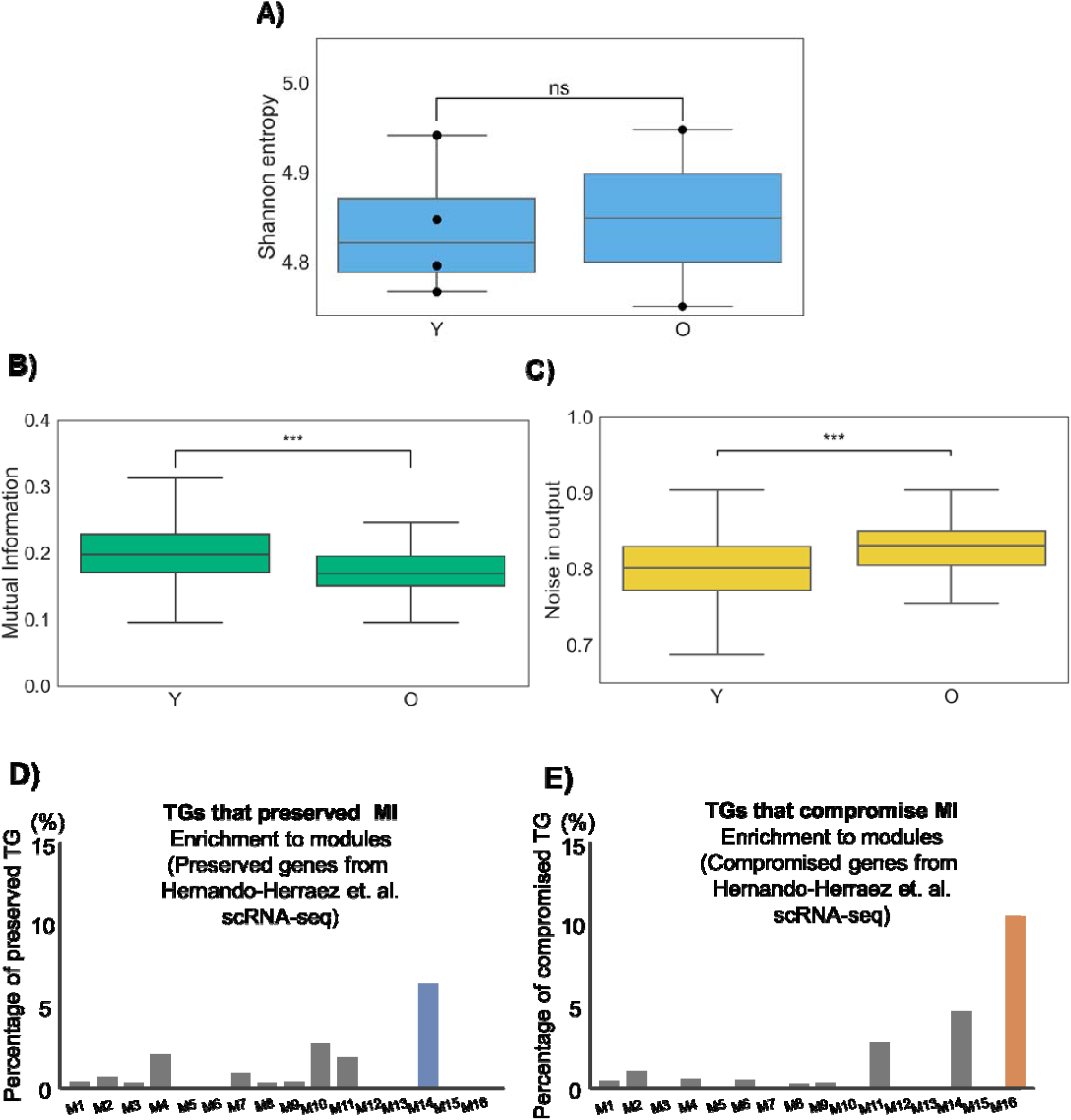
Reproduction of analysis with secondary dataset. **(A)** Shannon Entropy of mean gene expression per animal increases with age using single-cell RNA seq generated by Hernando-Herraez et. al. (*3*) (Number of cells in young = 253 and old = 124; Number of animals, all males: Young(Y) n=4, Old(O) n=2). The boxplot with overlaid individual data points indicates median and interquartile range. The increase is not statistically significant. **(B)** Mutual information decreases with age. **(C)** Noise in output increases with age. [**B, C**: Both mutual information and noise in the output (H(TG|TF)) were normalized by mean of entropy of input and output, as described in the methods. The boxplots represent median and interquartile range across age. *** represent p<0.01 for Bonferroni post-hoc performed after non-parametric Kruskal Wallis test.] **(D)**Similar to Figure 4C, Module 14 (blue bar) of the WGCNA network was enriched for ∼7% of TGs that preserved MI in the Hernando-Herraez et. al. **(E)** Similar to Figure 4C, Module 16 (orange bar) of the WGCNA network was enriched for ∼12% of TGs that preserved MI in the Hernando-Herraez et. al.

