## Supplementary material for "The fidelity of genetic information transfer with aging skeletal muscle segregates according to biological processes": Supplemental_2025_02_11.docx

Supplemental File

### Four-state model channel capacity calculations

Calculating mutual information and channel capacity requires knowledge of the underlying joint probability distribution of the gene pairs. These distributions are non-trivial. Therefore, to calculate mutual information and channel capacity, one needs to rely on algorithms to estimate probability densities from finite samples, which is its own field of research *(35, 36, 37)*. With the goal of calculating mutual information and channel capacity analytically, we coarsened the data into four distinct states, according to whether the target gene (Y) is on (1) or off (0) while the transcription factor (X) is on (1) or off (0). **Supplemental Figure 2** below shows an example of how the data from a gene pair is coarsened into four states.

### Mutual Information and Channel Capacity Derivations

With the data in the four-state model, the joint probability distribution can be easily calculated by counting the number of points in each respective state and then dividing by the total number of data points. Let these probabilities be represented by *a*, *b*, *c*, and *d*.

$$p\left( x,y \right)=\left\{ \begin{aligned} a if x=0, y=0 \\ b if x=1, y=0 \\ c if x=0, y=1 \\ d if x=1, y=1 \end{aligned} \right.$$

(1)

From Equation 1 the marginals *p*(*x*) and *p*(*y*) can be calculated along with the conditional

distribution *p*(*y*|*x*).

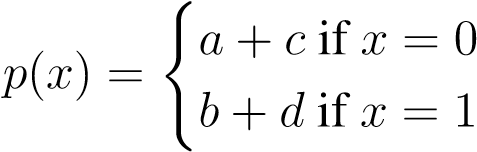
 (2)

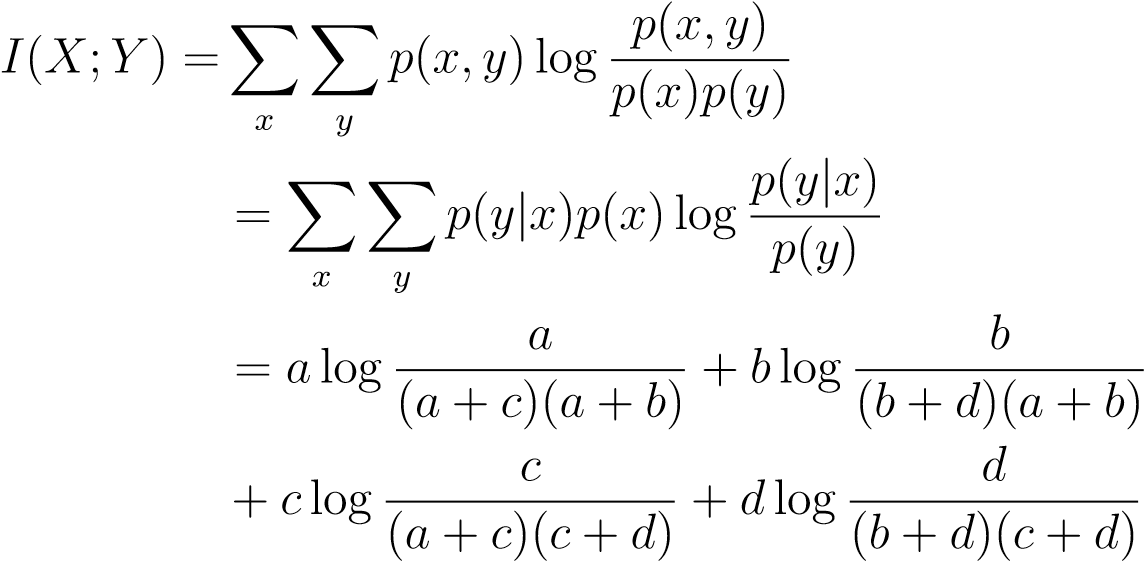

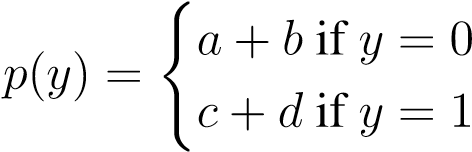
(3)

(4)

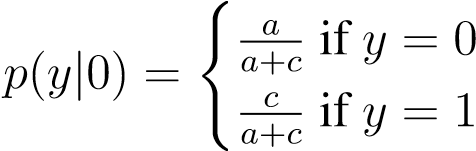
 (5)	(6)
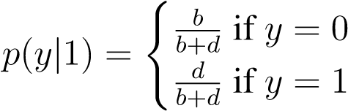

With the joint and marginal distributions in hand, the mutual information can be calculated.

To find the channel capacity, the maximum of *I* (*X*; *Y*) over all input distributions *p*(*x*) must be calculated while keeping the conditional distribution *p*(*y*|*x*) fixed. Let *α* = *a*+*c*,

*γ* = *c/*(*a*+*c*), and *β* = *b/*(*b* + *d*). Now Equations 2, 4, and 5 become

$p(x) = \left\{ \begin{aligned} \alpha if x=0 \\ 1- \alpha if x=1 \end{aligned} \right.$	(7)

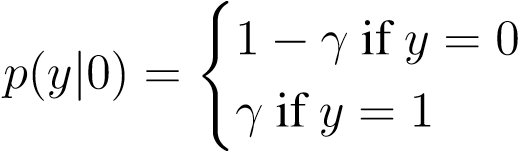
(8)

$p(y|1) = \left\{ \begin{aligned} \beta if y=0 \\ 1- \beta if y=1 \end{aligned} \right.$	(9)

From Equations 2, 4, and 5, we see that to maximize *I* (*X*; *Y*) over all input distributions *p*(*x*) while keeping the conditional distribution *p*(*y*|*x*) fixed, we need to optimize *I*(*α,β,γ*) with respect to *α* while keeping *γ* and *β* constant. Substituting *α*, *β*, and *γ* into Equation 6 we get:

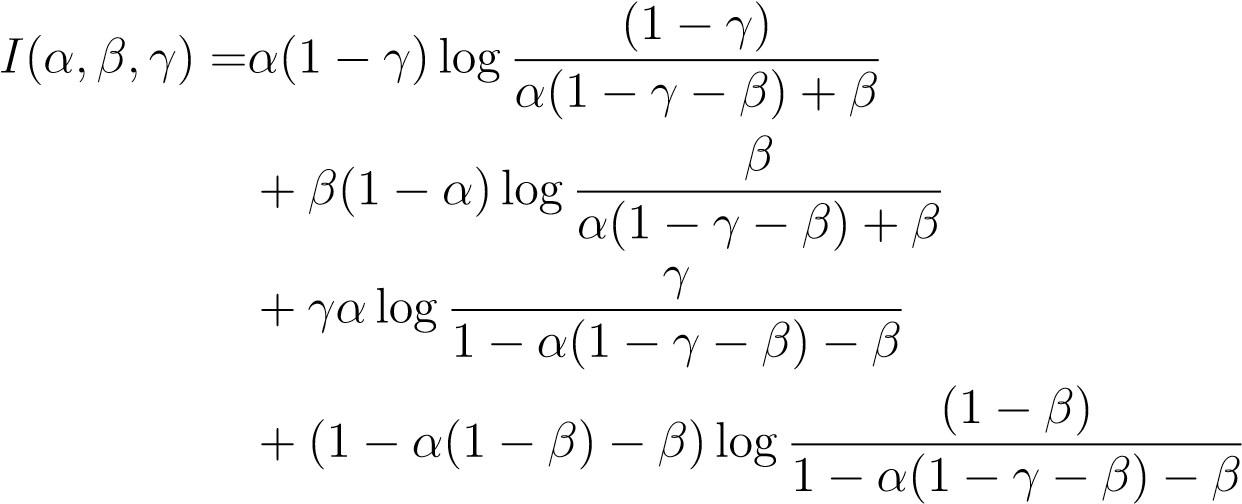

(10)

To simplify notation, let:

*χ* = *α* (1 − *β* − *γ*) + *β*	(11)

Now to optimize *I*(*α, β, γ*), we take its partial derivative with respect to *α*, set it equal to 0, and solve for *α*:

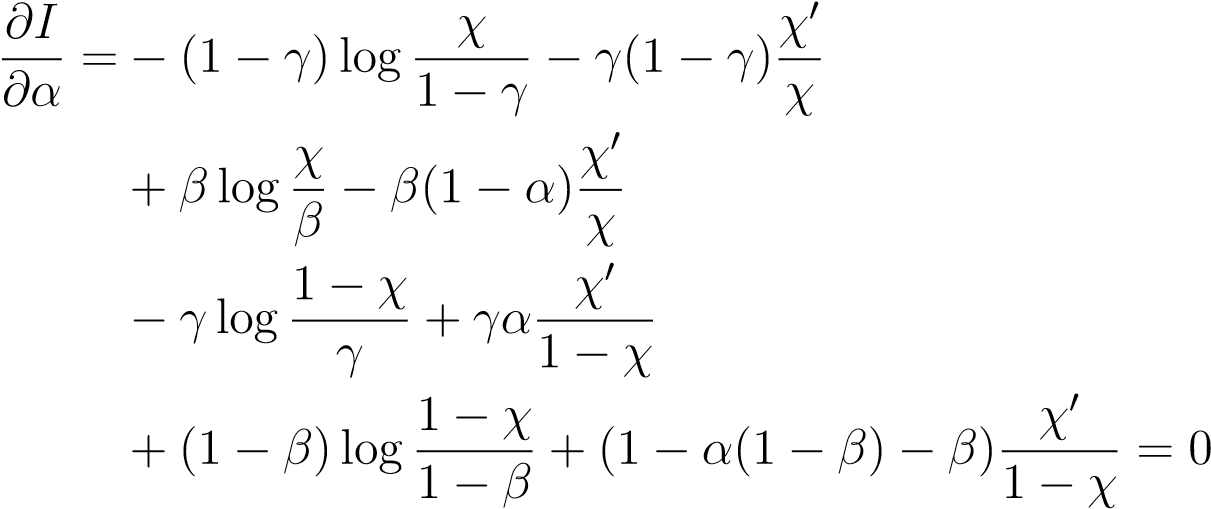
 (12)

*.*

The non-log terms in Equation 12 sum to 0. Collecting the log terms together we have:

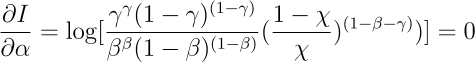
*.*	(13)

This means that:

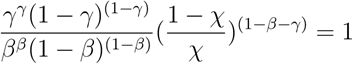
(14)

or

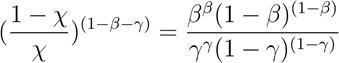
(15)

Let:

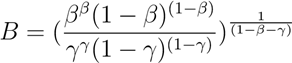
(16)

Now Equation 15 becomes:

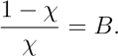
 (17)

Solving for *χ* we find the optimal *χ* value:

*.*
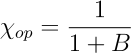
(18)

Plugging *χ_op_* into Equation 11 gives the value of *α* that optimizes *I*:

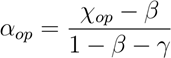
(19)

| The channel capacity is therefore: |  |
| --- | --- |
| *C* = *I* (*α_op_, β, γ*) | (20) |

### Mean Entropy

The mean entropy and mean optimal entropy are used to normalize the mutual information and channel capacity respectively. Entropy is defined as:

*H* = − *∑ p* log *p.*	(21)

The marginal distributions (Equations 2 and 3) can be written in terms of *α* and *χ:*

$p(x) = \left\{ \begin{aligned} \alpha if x=0 \\ 1- \alpha if x=1 \end{aligned} \right.$	(22)

$p(y) = \left\{ \begin{aligned} \chi if y=0 \\ 1- \chi if y=1 \end{aligned} \right.$	(23)

This gives the mean entropy and mean optimal entropy as:

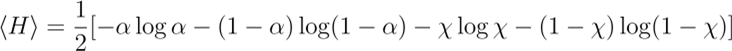
 (24)

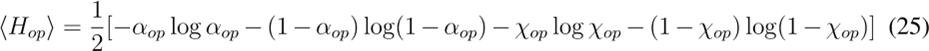

### Range and variability as they change with channel capacity

In this section we define range and variability. Both quantities have statistics involving the conditional probability p(TG=y|TF=x) (also known as the transfer function). The conditional probability is a family of distributions, Figure 2A, represents a way to show their statistics together in a single plot. The blue line represents the conditional averages while the red lines represent the conditional standard deviations. Both are defined below in Equations 26/29 and 28/31, respectively. The average of these conditional standard deviations is known as the variability. Range is defined below in Equation 32. The difference between maximum and minimum conditional average is the range.

We sought to understand why variability decreased as observed in Figure 2E. Recall that our input *X* (transcription factor) can either be a 0 or a 1 and our output *Y* (target gene) can also be either a 0 or a 1.

Using the definitions of average and standard deviation:

For *x* = 0, we have:

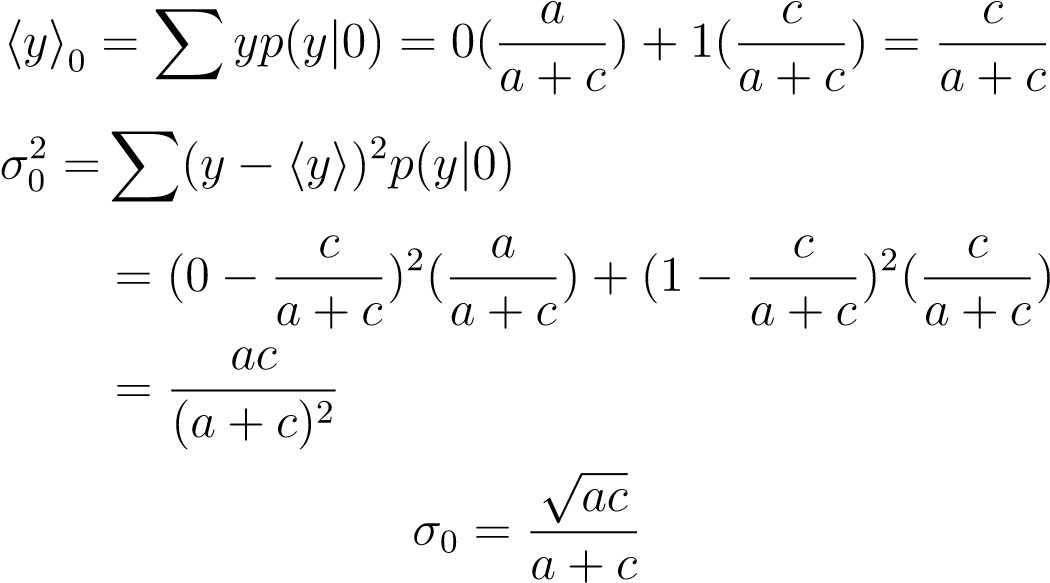
 (26)

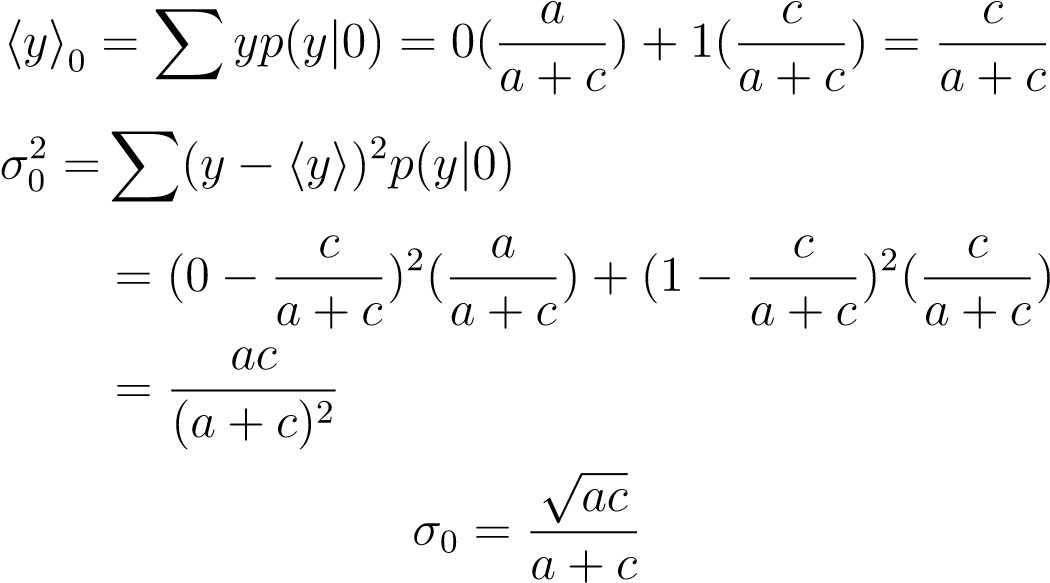

(27)

(28)

For *x* = 1, we have

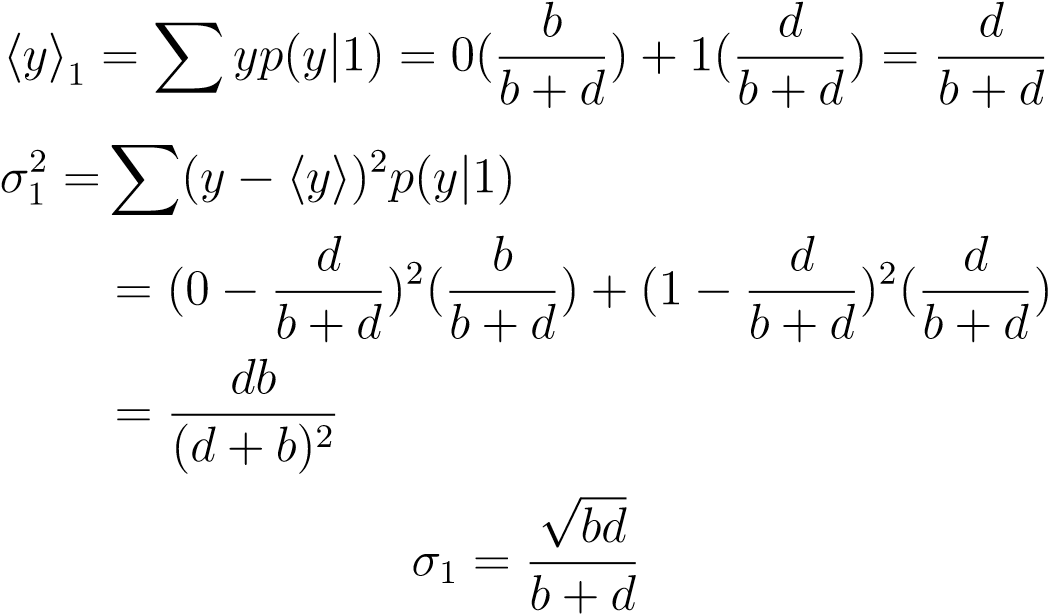
(29)

(30)

(31)

To see how the transfer function changes with time, we compare the ranges and standard deviations (variabilities) between young and aged. The range of the transfer function is defined to be:

*R* = |⟨*y*⟩_1_ − ⟨*y*⟩_0_|. (32)

Table 1 below shows whether the range and standard deviations increase or decrease with respect to age for all the genes.

|  | Categories | All  # genes (%) |
| --- | --- | --- |
| **1.** | Total Pairs | 3021 |
| **2.** | Decreasing Range | 1981 (.66) |
| **3.** | Increasing Range | 1040 (.34) |
| **4.** | Decreasing *σ*_0_ | 2571 (.85) |
| **5.** | Increasing *σ*_0_ | 450 (.15) |
| **6.** | Decreasing *σ*_1_ | 2409 (.80) |
| **7.** | Increasing *σ*_1_ | 612 (.20) |

Table 1: Transfer function statistics

The differences in the range and standard deviations are defined to be:

*Rdiff* = *Raged* − *Ryoung*	(33)

*σ_0,diff_* = *σ_0,aged_* – *σ_0,young_*	(34a)

*σ1,diff* = *σ1,aged* – *σ1,young*	(34b)

Table 2 below shows the averages of the differences in the range and standard deviations between young and old.

|  | All |
| --- | --- |
| Average *R_diff_* | -0.021 |
| Average *σ*0*,diff* | -0.076 |
| Average *σ*1*,diff* | -0.070 |

Table 2: Average differences in ranges and standard deviations

As can be seen from Table 1, the conditional standard deviation of the mean expression level decreases with age most of the time for both high (*x* = 1) and low (*x* = 0) transcription factor. This is attributed to the mean expression level, ⟨*y*⟩, decreasing with age. From equations 26*/*29 and 28*/*31, it can be seen that the conditional standard deviation can be written as:

*σ_0_* =	√⟨*y*⟩*_0_* (1 − ⟨*y*⟩*_0_*)	(35a)

*σ_1_* =	√⟨*y*⟩*_1_* (1 − ⟨*y*⟩*_1_*)	(35b)

This is a concave function (upside-down parabola opening downward) that has a maximum at *<y>* = 0*.*5 and minima at *<y>* = 0*,*1. This equation tells us that if the mean expression level, <y>*_young_* is less than 0*.*5 and if this mean decreases with age (<y>*_young_* greater than <y>*_old_* ) then the standard deviation (*σ*) necessarily has to decrease with age. Table 3 below shows the percentage of gene pairs (from Table 1, rows 4 and 6 with a decreasing standard deviation from young to old) that have <y>*_young_* less than 0*.*5 and whose mean decreases with age (<y>*_young_* greater than <y>*_old_* ).

|  | All |
| --- | --- |
| *x=0* | 0.93 |
| *x=1* | 0.88 |

Table 3: Percentage of gene pairs that have decreasing standard deviation with age, ⟨*y*⟩*_young_* less than 0*.*5, and whose mean decreases with age. This can happen for either x=0 or x=1.

### Four-state model with a non-zero threshold

The zero/nonzero coarse-graining in the four-state model is a large reduction from the original data. To see how much our conclusions depend on this strong reduction, we calculated the average mutual information and average channel capacity across gene pairs using FACS-Smartseq2 TMS limb muscle data while applying different non-zero threshold levels. Similarly defined as before, the four distinct states are now whether the target gene (Y) is larger (1) or smaller (0) than the threshold value while the transcription factor (X) is larger (1) or smaller (0) than the threshold value. The results are shown below in **Supplemental Figure 3**.

For reasonable non-zero threshold values (See Figure 1A for mean expression levels across cells and Supplemental Figure 2A for an example of expression levels in a given TF-TG pair), the mutual information decreases from 3 months to 24 months (**Supplemental Figure 3**). While the values of mutual information monotonically decrease with the modified thresholds, the decreasing trend with respect to age persists until extreme thresholding pushes all data into state 1 and the mutual information goes to zero. Channel capacity follows a similar trend. For a given threshold, channel capacity almost always decreases from 3 months to 24 months (**Supplemental Figure 3**). It is interesting to note that the capacity increases with threshold before ultimately decreasing. This increase is not well understood and is a question for future work.
